# SIRT5-dependent regulation of ASL controls arginine metabolism and T cell function

**DOI:** 10.64898/2026.06.24.734337

**Authors:** Kim Han, Rachael J. Klein, Kaiyuan Wu, Anna Chiara Russo, Allison M. Meadows, Thomas C. Recupero, Rahul Sharma, Anand K. Gupta, Nathan T. Brandes, Benjamin Schwarz, Michael N. Sack

## Abstract

NAD⁺ supplementation blunts Th1 and Th17 inflammation, in part, through arginine metabolism-dependent regulation of mitochondrial energetics, redox balance and signal transduction. Whether the NAD^+^-dependent sirtuin deacylases contribute to this regulation is unknown. Here, we show that both SIRT1 and SIRT5 transcript levels are induced in CD4^+^ T cells in human participants following oral supplementation of the NAD^+^ precursor nicotinamide riboside (NR). Among the sirtuin family members, SIRT5 rather than SIRT1 emerged as the predominant regulator of arginine and fumarate metabolism. Genetic depletion or pharmacologic inhibition of SIRT5 attenuated NR-mediated increases in arginine and fumarate and abolished the anti-inflammatory -effects of NR on Th1 and Th17 cytokine production. In contrast, the responses to exogenous arginine or citrulline supplementation were preserved, indicating that SIRT5 functions upstream of arginine biosynthesis. Metabolomic profiling further demonstrated that SIRT5 is required for NR-induced remodeling of the arginine biosynthetic pathway. Mechanistically, SIRT5 physically interacted with arginosuccinate lyase (ASL), promoted ASL-dependent arginine accumulation, and regulated ASL post-translational acylation, including glutarylation and malonylation. Loss of SIRT5 disrupted NR-mediated redox homeostasis, antioxidant gene expression, and cytokine suppression. Collectively, these findings identify SIRT5 as a critical mediator of NAD⁺ precursor-induced metabolic remodeling that links ASL-dependent arginine metabolism to redox balance and effector function in human CD4⁺ T cells.

## 1. Introduction

Nicotinamide adenine dinucleotide (NAD^+^) levels modulate multiple cellular functions via its role as a coenzyme for redox reactions, as a cofactor for multiple non-redox enzymes including sirtuins and for RNA NADylation.[1] These diverse roles of NAD^+^ introduce complexity in exploring its biological function. Concurrently vitamin B_3_ analogue supplementation, as precursors to increase intracellular NAD^+^ levels, are increasingly being employed as potential therapeutic interventions.[2] This has stimulated efforts to better understand the mechanisms through which NAD⁺ regulates human physiology and pathology. However, a recent review questioned the utility of NAD^+^ supplementation in humans, except for a possible regulatory role in immune modulation.[3] In pilot human studies, NAD^+^ supplementation has shown to confer anti-inflammatory[4–6] and potential anti-tumorigenic effects,[7] although the mechanisms of action remain incompletely characterized.[8]

Our laboratory has undertaken numerous studies to explore the mechanisms of action of the oral NAD^+^ precursor nicotinamide riboside (NR) in healthy individuals, and in participants with psoriasis and systemic lupus erythematosus (SLE)[9, 10] and an ongoing placebo-controlled study in SLE (NCT06032923). To date, these studies highlight that increasing whole blood NAD^+^ levels confer effects on metabolic pathways, signal transduction, organelle homeostasis and bioenergetic and redox control in myeloid and CD4^+^ T cells and more recently in non-immune cells including skin fibroblasts.

Historically, we have shown that the NAD^+^-dependent mitochondrial enriched sirtuin enzyme SIRT3 plays a role in attenuating the NLRP3 inflammasome in myeloid cells.[11, 12] This and other sirtuin enzymes have also been shown to confer a multitude of anti-inflammatory effects in different myeloid cell lineages.[8] In parallel, studies have begun to explore the role of SIRT1 and SIRT3 in CD8^+^ and B cell immunity.[8] The roles of these enzymes in CD4^+^ T cells are, however, less well established. At the same time, SIRT5 has been implicated in the control of urea cycle enzyme activities, and we have found that NR augments arginine biosynthesis as a driver of its Th1 and Th17 anti-inflammatory effects. Taken together, these findings suggest that sirtuin biology may contribute towards the immune modulatory effects of NR on CD4^+^ T cells.

In this study, we systematically screened the effect of the different sirtuin depletion on arginine and fumarate metabolism in human CD4^+^ cells. Among the seven sirtuin family members, only SIRT5 depletion consistently blunted the levels of these metabolic intermediates in the arginine biosynthesis pathway. Subsequent analysis showed that the depletion of SIRT5 attenuated the NR supplementation role in augmenting arginine biosynthesis and on dampening Th1 and Th17 immune responsiveness. Steady-state metabolomic analysis in CD4^+^ T cells further supports the concept that SIRT5 is an essential mediator of NR-induced arginine biosynthesis. Furthermore, we identify arginosuccinate lyase (ASL) as a SIRT5-interacting protein and show that SIRT5 regulates ASL acylation, with deglutarylation emerging as the predominant modification associated with NR-induced arginine biosynthesis.

## 2. Materials and methods

### 2.1. Study Approval and Trial Registration

Human samples were obtained from participants enrolled in clinical protocols approved by the Institutional Review Board of the NIH Clinical Center. All participants provided written informed consent. Clinical trial registration numbers were NCT02719899, NCT02812238, and NCT01143454. Additional samples were also obtained through the NIH Clinical Center Blood Bank under protocol NCT00001846 (ClinicalTrials.gov).

### 2.2. Cell Isolation and Culture

Primary human CD4⁺ T cells were isolated from peripheral blood mononuclear cells (PBMCs) using a negative selection kit (Miltenyi Biotec). Cells were cultured in RPMI 1640 medium (Invitrogen) supplemented with 10% heat-inactivated fetal bovine serum (FBS; Invitrogen), 1% Antibiotic-Antimycotic solution (Thermo Fisher Scientific), and 25 mM HEPES (pH 7.4) at 37°C in a humidified atmosphere containing 5% CO₂. Cells were maintained under non-polarizing (Th0) conditions. For T cell receptor (TCR) activation, culture plates were pre-coated with anti-human CD3 antibody (clone OKT3, BioLegend: 5 μg/ml) and anti-human CD28 antibody (clone CD28.2, BioLegend: 10 μg/ml) was added to the culture medium at the time of stimulation.

### 2.3. Plasmids and Transfection

The human ASL expression plasmid (ASL-Flag) was provided by Dr. Shi-Min Zhao (Fudan University). A full-length wild-type human ASL construct with a C-terminal Flag tag and human SIRT5 (HA-tagged or Myc-tagged) were obtained from Sino Biological and Addgene, respectively. Site-directed mutagenesis (K69R, K288R, and double mutant) was performed using the Q5 Site-Directed Mutagenesis Kit (New England Biolabs). Plasmids were transfected using PolyJet transfection reagent (SignaGen Laboratories). HA-tagged SIRT5 was used for co-immunoprecipitation studies, whereas Myc-tagged SIRT5 was used for ASL acylation analyses.

### 2.4. Inhibitors, siRNAs, and Lentiviruses

Cells were treated with nicotinamide riboside (NR; ChromaDex) at 0.5 mM for 72 hours as indicated. Where indicated, L-arginine (Sigma-Aldrich) was supplemented at 5 mM for 72 hours. SIRT5 inhibition was achieved using small-molecule inhibitors NRD167 (Selleckchem) or MC3482 (MedChem Express) at 10 μM for 24 hours. For knockdown experiments, SMARTpool Accell siRNAs targeting ASL and sirtuins (Horizon Discovery) were used. Cells were treated with Accell siRNA in Accell siRNA Delivery Media (Horizon Discovery) according to the manufacturer’s instructions. Cells (4-5 × 10⁶) were incubated with 1.5 μM siRNA and subsequently activated with plate-coated anti-CD3 (BioLegend) and anti-CD28 (BioLegend) antibodies for 72 hours prior to downstream assays. For lentivirus production, pLenti vector, pLenti-SIRT5 with Myc tag, or pLenti-ASL with Flag tag (OriGene Technologies), together with packaging plasmids psPAX2 and pMD2.G (Addgene), were transfected into HEK293T cells (ATCC) using PolyJet transfection reagent (SignaGen Laboratories). Supernatants were collected 48 hours post-transfection, filtered through a 0.45 μm syringe filter, and concentrated by ultracentrifugation (Beckman Coulter) at 25,000 rpm for 2 hours. For transduction, concentrated virus was added to CD4⁺ T cells in the presence of DEAE-dextran (Sigma-Aldrich, 8 μg/mL) and LentiBOOST Transduction Enhancer (Mayflower Bioscience) according to the manufacturer’s instructions.

### 2.5. Quantitative RT-PCR (qPCR)

Total RNA was extracted using the NucleoSpin RNA kit (Macherey-Nagel). cDNA was synthesized using the SuperScript III First-Strand Synthesis System for RT-PCR (Thermo Fisher Scientific) according to the manufacturer’s instructions. Quantitative real-time PCR was performed using FastStart Universal SYBR Green Master Mix (Roche) on a LightCycler 96 system (Roche). Target gene expression was analyzed using validated gene-specific QuantiTect primers (Qiagen) as listed in Supplemental Table 1. Relative gene expression was calculated using the comparative Ct (2^−ΔΔCt) method after normalization to housekeeping genes (18S rRNA or β-actin).

### 2.6. Metabolite and Cytokine Measurements

Intracellular metabolites were extracted from cells and normalized to protein content using a BCA assay (Thermo Fisher Scientific). L-arginine levels were quantified using a commercially available ELISA kit (Immusmol), and fumarate levels were measured using a fumarate assay kit (Sigma-Aldrich). Cell culture supernatants were collected, centrifuged to remove debris, and cytokine levels (IFNγ, IL-17, and IL-5) were measured using ELISA kits (R&D Systems). Cytokine concentrations were quantified using standard curves and normalized to cell number (CyQUANT Cell Proliferation Assay, Invitrogen) or total protein content where indicated.

### 2.7. Reactive Oxygen Species (ROS) Assays

Intracellular and mitochondrial reactive oxygen species (ROS) were measured using both flow cytometry and microplate reader–based assays. CD4⁺ T cells (4-8 × 10⁵ cells per well) were incubated with 2’,7’-dichlorofluorescin diacetate (DCFDA; Thermo Fisher Scientific or Sigma-Aldrich) at 10 μM or MitoSOX™ Red mitochondrial superoxide indicator (Thermo Fisher Scientific/Invitrogen) at 10 μM in HBSS for 30 minutes at 37°C. Cells were subsequently washed with HBSS prior to analysis. DCFDA fluorescence (total ROS) was detected at Ex/Em = 485/535 nm (plate reader) or using a 488 nm excitation laser (flow cytometry). MitoSOX Red fluorescence (mitochondrial superoxide) was detected at Ex/Em = 510/580 nm. Results were normalized to total protein content using the Pierce BCA Protein Assay (Thermo Fisher Scientific) and to cell number using the CyQUANT Cell Proliferation Assay (Invitrogen).

### 2.8. Metabolomics and Data Analysis

CD4⁺ T cells were transfected with non-targeting control (N.C.) or SIRT5-targeting siRNA and subsequently treated with NR (0.5 mM) or L-arginine (5 mM) for 72 hours as indicated. Cells were activated with plate-coated anti-CD3 and soluble anti-CD28 for 72 hours. Cell pellets were collected, snap-frozen in liquid nitrogen, and subjected to metabolomic analysis by liquid chromatography–mass spectrometry (LC-MS). Metabolite abundance was normalized to total protein content. Multivariate analysis was performed using SIMCA software (version 15, Umetrics) with orthogonal partial least squares discriminant analysis (OPLS-DA) to identify variable importance in projection (VIP) metabolites distinguishing treatment groups. Pathway enrichment analysis of VIP metabolites was conducted using MetaboAnalyst 5.0. Targeted metabolomics was performed using LC-MS/MS. Samples were extracted using a methanol-based extraction method and analyzed using a Vanquish™ UHPLC system coupled to a TSQ Quantiva™ mass spectrometer (Thermo Fisher Scientific). Reverse-phase chromatography was performed using a C18 column under standard conditions. Electrospray ionization was operated in both positive and negative ionization modes. Data acquisition and processing were performed using Xcalibur software (version 2.2, Thermo Fisher Scientific). Peak intensities were normalized to total signal intensity. Statistical analyses were performed using paired t-tests, with false discovery rate (FDR) correction thresholds set at 10-20% as indicated. Data visualization included volcano plots, heatmaps, and hierarchical clustering generated using GraphPad Prism (version 11).

### 2.9. MetLip

Tributylamine and all synthetic molecular references were purchased from Millipore Sigma. LCMS grade water, methanol, isopropanol and acetic acid were purchased through Fisher Scientific. Aqueous metabolites were analyzed using a combination of two analytical methods with opposing ionization polarities.[13] Both methodologies utilized a LD40 XR UHPLC (Shimadzu Co.) system for separation and a 6500+ QTrap mass spectrometer (AB Sciex Pte. Ltd.) for detection. Negative mode samples were separated on a Waters™ Atlantis T3 column (100Å, 3 µm, 3 mm X 100 mm) with a gradient from 5 mM tributylamine, 5 mM acetic acid in 2% isopropanol, 5% methanol, 93% water (v/v) to 100% isopropanol over 5 mins. Two distinct multiple reaction monitoring (MRM) pairs in negative mode were used for each metabolite for improved identification confidence where available. Positive mode method samples were separated across a Phenomenex Kinetex F5 column (100 Å, 2.6 µm, 100 x 2.1 mm) using a gradient from 0.1 % formic acid in water to 0.1 % formic acid in acetonitrile over 5 minutes. All signals were integrated using SciexOS 3.1 (AB Sciex Pte. Ltd.). Signals were inspected visually for presence and signals with greater than 50% missing values were discarded. Any remaining missing values were replaced with the minimum detected value within the dataset. Metabolites with multiple MRMs were quantified with the higher signal to noise MRM. Filtered datasets of the negative mode aqueous metabolites were total sum normalized after initial filtering. The positive mode was scaled and combined with the normalized negative mode dataset via common signals for Threonine. A Benjamini-Hochberg method for correction for multiple comparisons was imposed where indicated.

### 2.10. Immunoprecipitation and Immunoblotting

HEK293 cells were lysed in RIPA buffer (Boston BioProducts) supplemented with protease inhibitor cocktail (Roche) and phosphatase inhibitors (Pierce). For immunoprecipitation, clarified lysates were incubated overnight at 4°C with anti-Flag antibody (Sigma-Aldrich) or anti-SIRT5 antibody (Cell Signaling Technology). Where indicated, Flag-tagged proteins were immunoprecipitated using magnetic beads conjugated with anti-Flag M2 or anti-HA (Sigma-Aldrich) and eluted with Flag peptide (Sigma-Aldrich) according to the manufacturer’s instructions. Alternatively, immune complexes were captured using magnetic Protein A/G beads (SignaGen Laboratories) and washed extensively. For immunoblotting, primary human CD4^+^ T cells or HEK293T cells were lysed as described above, and lysates were separated on NuPAGE 4-12% Bis-Tris protein gels (Thermo Fisher Scientific) and transferred to nitrocellulose membranes using the Trans-Blot Turbo Transfer System (Bio-Rad Laboratories). Membranes were blocked with Odyssey Blocking Buffer (Li-Cor Biosciences) and incubated with primary antibodies overnight at 4°C. Primary antibodies included ASL (Abcam or Proteintech), SIRT5 (Cell Signaling Technology), Flag (Sigma), Myc (Proteintech), HA (Roche), and β-Actin (Millipore) as a loading control. Membranes were then incubated with IRDye 800CW or IRDye 680RD-conjugated secondary antibodies (Li-Cor Biosciences) for 1 hour at room temperature. Immunoblots were scanned using an Odyssey DLx imaging system (Li-CorBiosciences). Protein band intensity was quantified using Image Studio software (version 5.2, Li-Cor) and normalized to β-Actin levels.

### 2.11. Quantification and Statistical Analysis

Statistical analyses were performed using GraphPad Prism (version 11). Data are presented as mean ± standard error of the mean (SEM) unless otherwise indicated. Comparisons between two groups were performed using paired or unpaired two-tailed Student’s t-tests as appropriate. For comparisons involving more than two groups, one-way analysis of variance (ANOVA) was used, followed by appropriate post hoc tests. Two-way ANOVA was applied when two independent variables were analyzed. A p-value < 0.05 was considered statistically significant. For metabolomics data, differential metabolite levels were assessed using paired t-tests with false discovery rate (FDR) correction. Adjusted p-values < 0.05 were considered significant. Metabolite changes were visualized as log₂ fold change in metabolic pathway maps, as well as in volcano plots and heatmaps. For all experiments, n represents the number of biological replicates and is indicated in the figure legends.

## 3. Results

### 3.1. Fasting and NAD boosting coordinately regulate sirtuin signaling and arginine metabolism

Our laboratory initially employed acute fasting as a human intervention to study CD4^+^ T cell immunometabolism and regulation.[14] As a component of this intervention we employed integrative lipidomic, metabolomic and bulk RNAseq bioinformatics analysis comparing out 24-hour fasting compared to 3 hour refeeding studies,[14–16] and identified by Ingenuity pathway analysis (IPA) that sirtuin signaling pathways and arginine biosynthesis pathways were both significantly enriched with canonical transcript and metabolite effects (Fig. 1A). In parallel, and to enable longer-term placebo-controlled human interventions, we switched to investigating nicotinamide riboside (NR) supplementation, which increases blood NAD^+^-levels, as a canonical biochemical responses to caloric restrictive interventions.[17] Interestingly, we identified that NR functions in part via augmentation of arginine biosynthesis.[9] The integration of both of these dietary interventions, coupled to the established role of sirtuin deacylase SIRT5 in the regulation of the urea cycle and macrophage polarization,[18, 19] raised the question as to whether sirtuin biology and arginine metabolism may be integrated in NR-mediated blunting of Th1 and Th17 immune responsiveness. The hypothesis we explore in this manuscript is whether NR supplementation activates SIRT5 to induce arginine biosynthesis via modulating CD4^+^ T Cell arginosuccinate lyase (ASL) to control their immunometabolism and immune responsiveness (Fig. 1B). To pursue this line of investigation we then quantified the relative expression of the sirtuin isoforms in naïve human Th0 cells. Interestingly, the transcript levels of *SIRT1* were the most abundant, with *SIRT3* and *4*, the least abundant with intermediate levels of *SIRT2/5/6* and *7* (Fig. S1A). We then assessed whether SIRT5 mRNA levels were modified by fasting, and here we showed its induction in the 24-hour fasted, compared to the refed state in CD4^+^ T cells (Fig. S1B). To begin to evaluate the effect of NR, we assayed the effect of NR supplementation on sirtuin transcript levels in primary human CD4^+^ T cells. Here, NR significantly increased the expression of *SIRT1*, *SIRT3* and *SIRT5* (Fig. 1C). As we had previously studied the in-vivo effect of NR on immune function in healthy participants (NCT02812238) we then quantified the effect of 1 week of oral NR supplementation, as described previously,[9] on sirtuin levels. Interestingly, here, oral NR supplementation significantly increased *SIRT1* and *SIRT5* but not *SIRT3* transcript levels (Fig. 1D). Interestingly, ex-vivo NR treatment did not induce expression of arginine biosynthesis enzymes and modestly reduced *ASL* transcript levels, suggesting that NR regulates arginine metabolism independently of transcriptional induction of this pathway (Fig. S1C).

**Figure 1.**
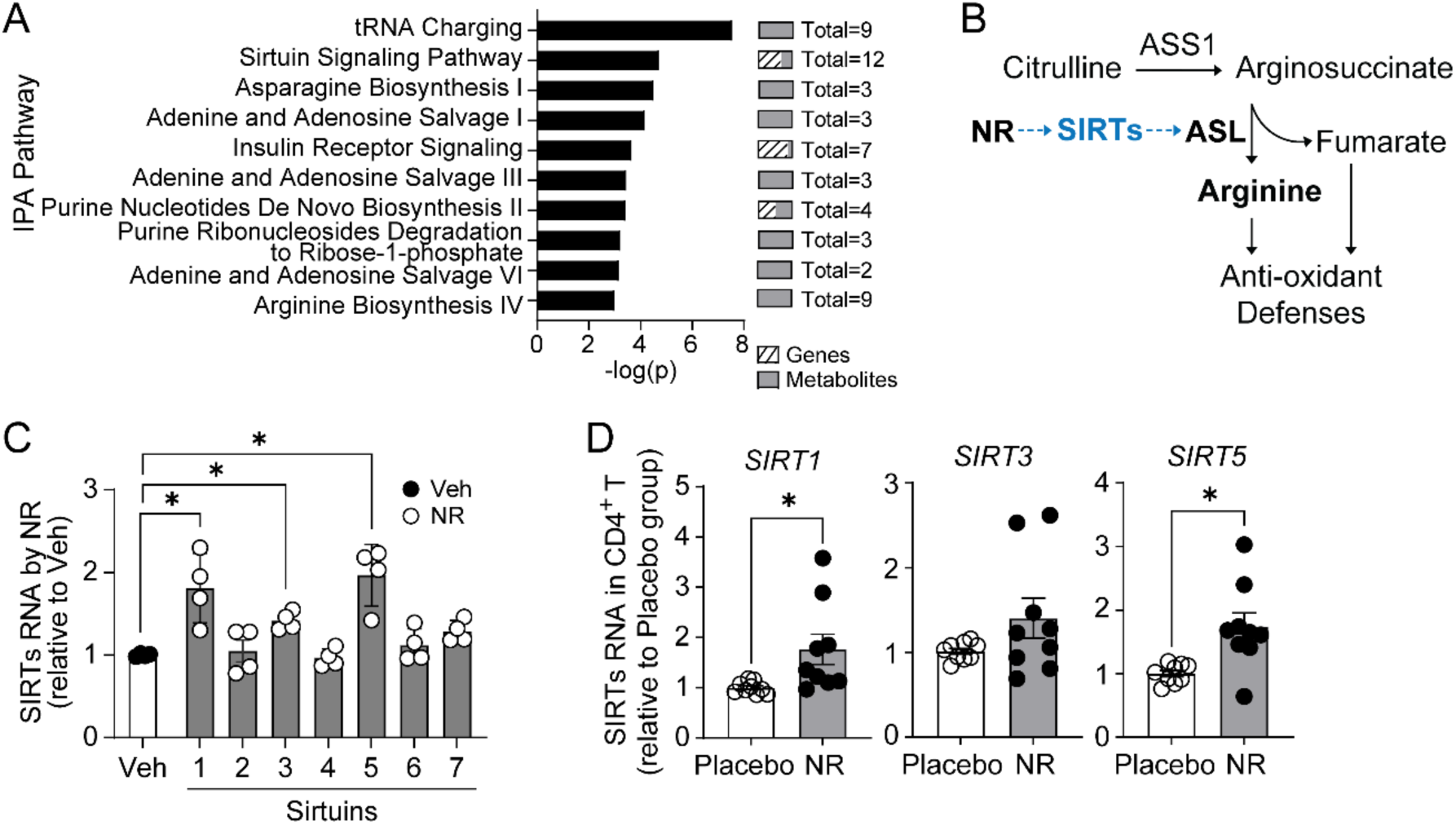
NR induces sirtuin signaling and arginine metabolism pathways in CD4⁺ T cells. (**A**) IPA analysis integrating lipidomics, metabolomics, and RNA-seq datasets identified enrichment of sirtuin signaling pathways in PBMCs following 24-hour fasting and 3-hour refeeding. Data were derived from a clinical study (ClinicalTrials.gov: NCT02719899). Differentially expressed genes, lipids, and metabolites used for this analysis are provided in Supplementary Dataset 1. (**B**) Schematic model linking NR-induced sirtuin activation to ASL-dependent arginine metabolism. (**C**) Relative RNA expression of sirtuin family members (*SIRT1-7*) in CD4⁺ T cells treated with NR compared to vehicle. (**D**) Relative expression of *SIRT1*, *SIRT3*, and *SIRT5* in CD4⁺ T cells from placebo and NR-treated groups derived from a clinical protocol (ClinicalTrials.gov: NCT02812238). Data in (C-D) are presented as mean ± SEM from independent biological replicates, with each dot representing an individual donor. Statistical significance in (C) was determined using one-way ANOVA with Šídák’s multiple comparisons test. Statistical significance in (D) was determined using an unpaired two-tailed Student’s *t*-test. \**p* < 0.05.

### 3.2. SIRT5 regulates arginine metabolism and cytokine production in CD4⁺ T cells

To further dissect out the putative roles of sirtuins in this regulation we employed siRNA depletion of the different isoforms in Th0 cells. siRNA was effective in reducing the transcript levels of all seven sirtuins with the most robust effects on *SIRT4/5* and *7* (Fig. S2A). We then assessed whether the knockdown of the individuals sirtuins affected CD4^+^ T cells arginine and fumarate levels. Interestingly, only the depletion of SIRT5 blunted the levels these metabolites, and this was to a similar extent to the siRNA targeting of *ASL* and *ASS1* (Figures S2B-D). To explore whether SIRT5 was necessary in the known NR effects on increasing arginine biosynthesis we employed siRNA targeting of SIRT5 in primary human Th0 cells. The depletion of SIRT5 attenuated the effect of NR to increase both arginine and fumarate levels (Figures 2A-B). In parallel the NR effect on diminishing the release of Th1 and Th17 cytokines, i.e. interferon gamma (IFNγ), interleukin 5 (IL-5) and IL-17, were also abolished following the depletion of SIRT5 (Fig. 2C). Interestingly, the direct supplementation of arginine abrogated the effect of SIRT5 depletion on the blunting of IFNγ, IL-5 and IL-17 levels (Fig. 2D). As NR had shown to attenuate the NLRP3 inflammasome in myeloid cells in a SIRT3 dependent manner,[11] we assayed whether SIRT3 was required for the effects of NR on Th1 and Th17 cytokine release. Although here, the knockdown of *SIRT3* did not attenuate the CD4^+^ T cell immune response in response to NR supplementation (Fig. S2E).

**Figure 2.**
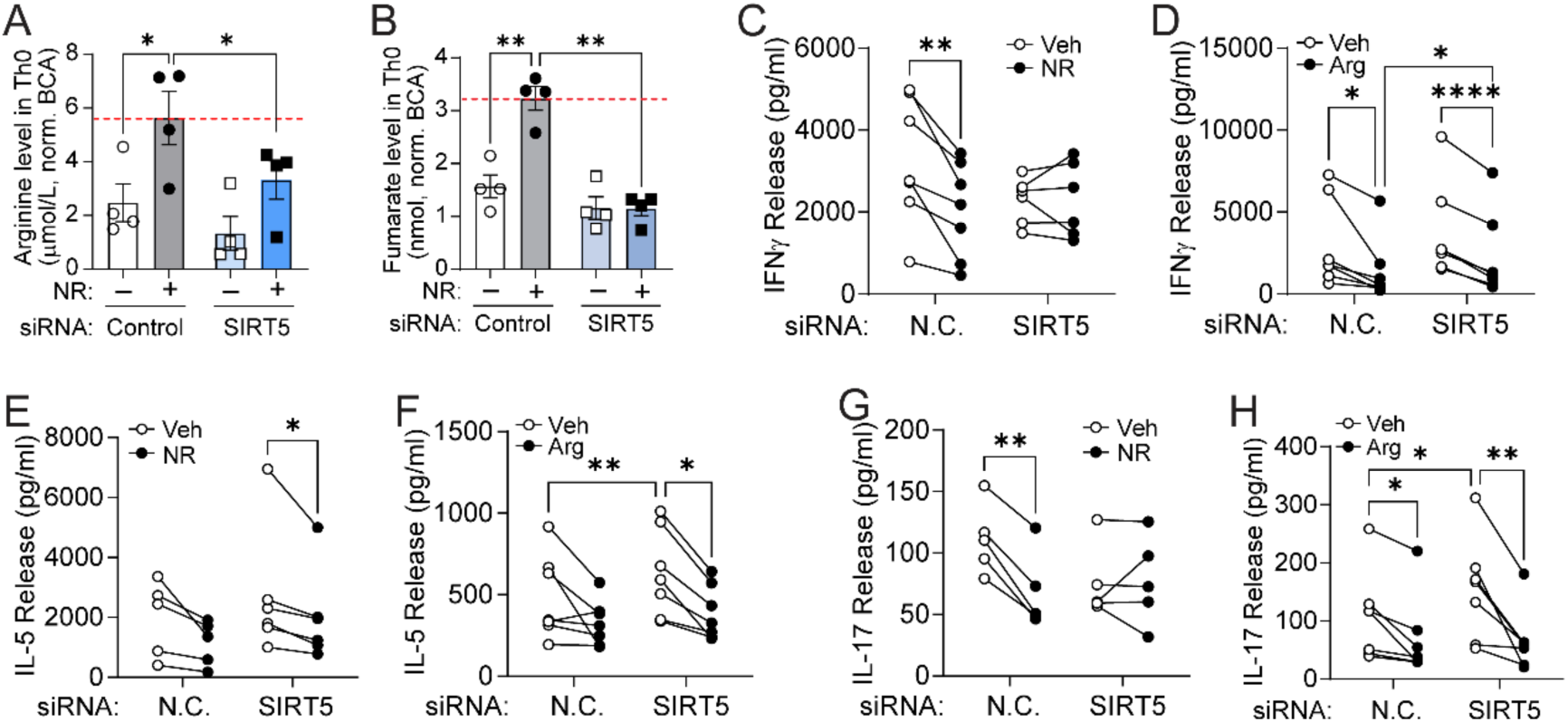
SIRT5 regulates arginine metabolism and cytokine production in CD4⁺ T cells. (**A**) Intracellular arginine levels in CD4⁺ T cells transfected with control or SIRT5 siRNA in the presence or absence of NR. (**B**) Intracellular fumarate levels under the same conditions. (**C-D**) IFNγ production in control (N.C.) and SIRT5 KD CD4⁺ T cells treated with NR (**C**) or arginine (**D**). (**E-F**) IL-5 production in control (N.C.) and SIRT5 KD CD4⁺ T cells treated with NR (**E**) or arginine (**F**). (**G-H**) IL-17A production in control (N.C.) and SIRT5 KD CD4⁺ T cells treated with NR (**G**) or arginine (**H**). Data are presented as mean ± SEM from independent biological replicates, with each dot representing an individual donor. Statistical significance was determined using two-way ANOVA followed by Šídák’s multiple comparisons test. \**p* < 0.05, \*\**p* < 0.01, \*\*\**p* < 0.001.

To validate that NR modulates arginine and fumarate synthesis rather than directly attenuating Th1/Th17 cytokine secretion, SIRT5 inhibitors NRD167[20] and MC3482[21] were employed as substitutes for the siRNA studies. Here too, the SIRT5 inhibitors reduced Th0 cell arginine and fumarate levels (Figures 3A-B) and attenuated the effect of NR on IFNγ and IL-17 release (Figures 3C-D). In parallel with the siRNA studies the SIRT5 inhibitors did not abolish the capacity of arginine supplementation itself to blunt the release of these cytokines (Figures 3E-F).

**Figure 3.**
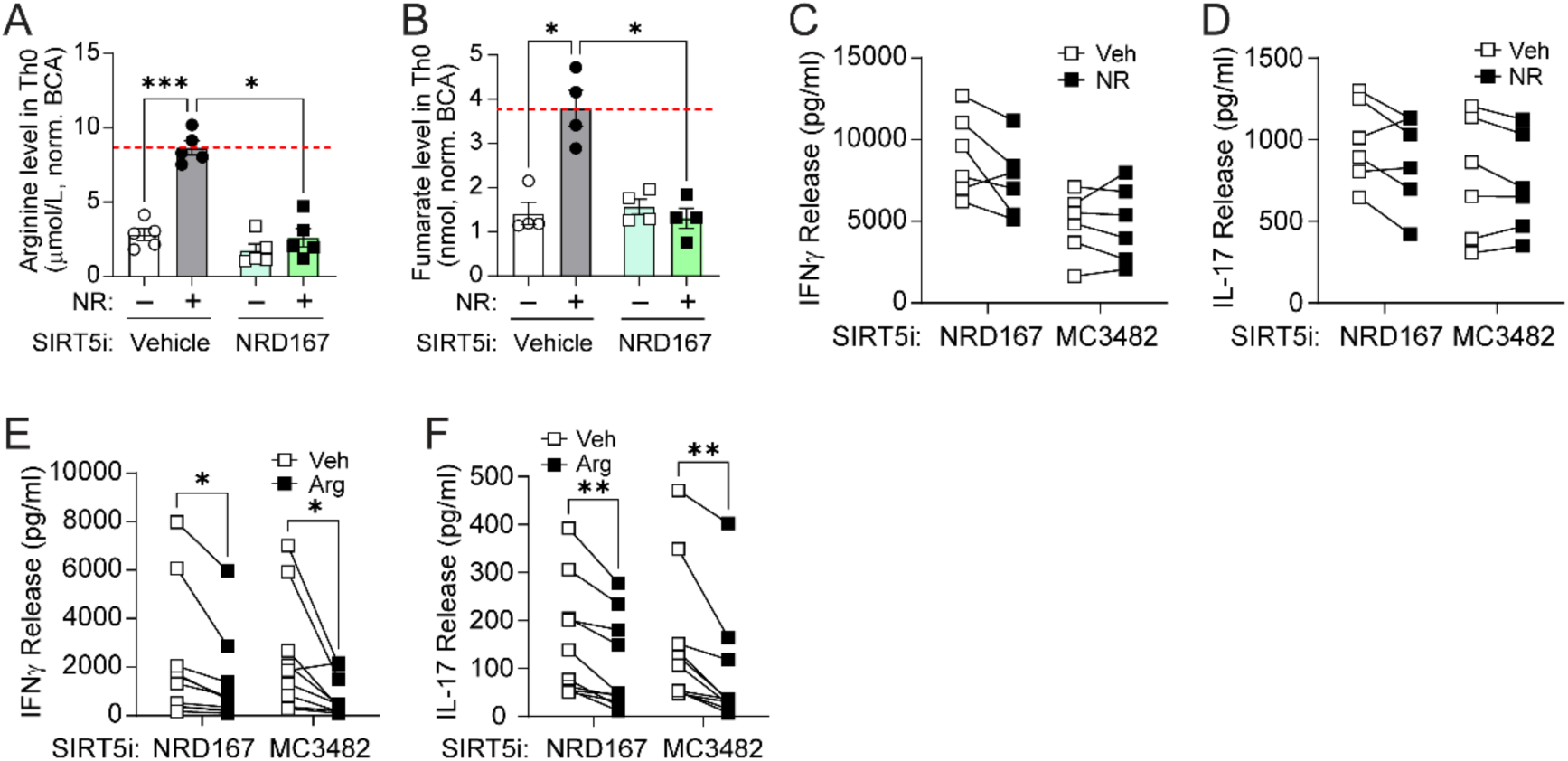
Pharmacologic inhibition of SIRT5 impairs NR-induced metabolic and functional responses. (**A**) Intracellular arginine levels in CD4⁺ T cells treated with NR in the presence or absence of the SIRT5 inhibitor NRD167. (**B**) Intracellular fumarate levels in CD4⁺ T cells treated with NR in the presence or absence of SIRT5 inhibitors (NRD167 or MC3482). (**C-D**) IFNγ and IL-17A production in CD4⁺ T cells treated with NR in the presence of SIRT5 inhibitors (NRD167 or MC3482). (**E-F**) IFNγ and IL-17A production in CD4⁺ T cells treated with arginine in the presence of SIRT5 inhibitors (NRD167 or MC3482). Data are presented as mean ± SEM from independent biological replicates, with each dot representing an individual donor. Statistical significance was determined using two-way ANOVA followed by Šídák’s multiple comparisons test. \**p* < 0.05, \*\**p* < 0.01, \*\*\**p* < 0.001.

### 3.3. SIRT5 regulates redox balance and antioxidant responses in CD4⁺ T cells

As the CD4^+^ T cell immunomodulatory effect of NR is dependent on reducing mitochondrial reactive oxygen species (ROS) levels,[9] we explored if SIRT5 is necessary in this signaling. Here we show that in response to SIRT5 depletion, the effect of NR on blunting whole-cell (DCFDA) and mitochondrial ROS (MitoSOX) fluorescence (Figures 4A-B). In contrast, and in parallel with the effects on cytokine release, SIRT5 was not necessary for the ameliorative effect of arginine supplementation on ROS levels (Figures 4A-B). This phenotype on mitochondrial ROS was replicated by SIRT5 inhibition with MC3482 (Fig. 4C) but not evident when measuring whole-cell ROS levels (Fig. 4D). Again here, SIRT5 inhibition did not blunt the effect of arginine supplementation on ROS levels (Figures 4C-D). In parallel with these findings both the genetic depletion of SIRT5 or its pharmacologic inhibition blunted transcript levels of canonical NR-responsive antioxidant gene transcript levels (Figures 4E-F).

**Figure 4.**
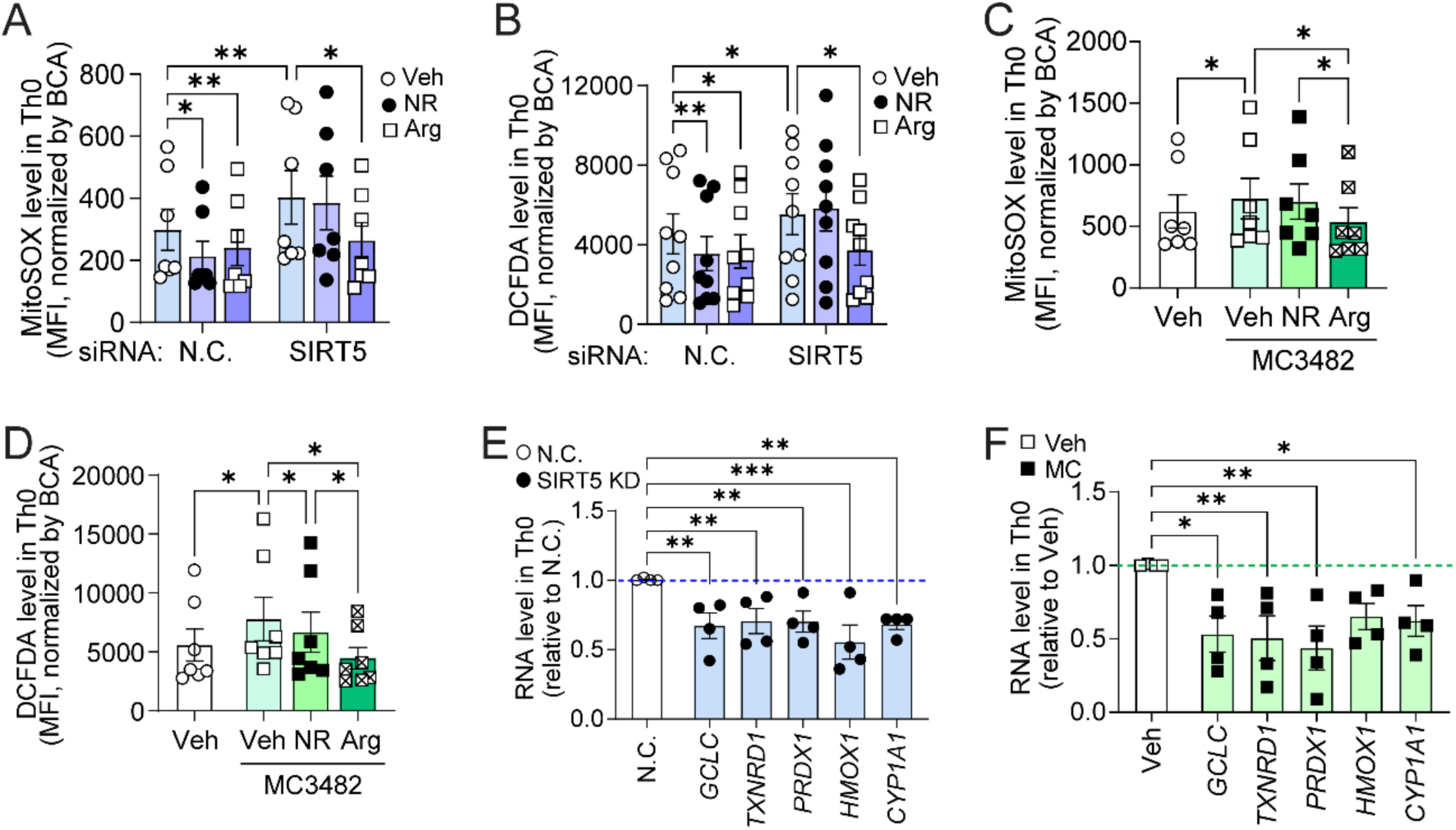
SIRT5 regulates redox balance and antioxidant responses in CD4⁺ T cells. (**A**) Mitochondrial superoxide levels measured by MitoSOX fluorescence under the same conditions. (**B**) Intracellular reactive oxygen species (ROS) levels measured by DCFDA fluorescence in control (N.C.) and SIRT5 KD CD4⁺ T cells treated with vehicle, NR, or arginine. (**C**) Mitochondrial superoxide levels measured by MitoSOX in CD4⁺ T cells treated with MC3482 under the same conditions. (**D**) Intracellular ROS levels measured by DCFDA in CD4⁺ T cells treated with the SIRT5 inhibitor MC3482 in the presence of vehicle, NR, or arginine. (**E)** Relative expression of antioxidant genes (GCLC, TXNRD1, PRDX1, HMOX1, and CYP1A1) in control (N.C.) and SIRT5 KD CD4⁺ T cells. (**F**) Relative expression of antioxidant genes in CD4⁺ T cells treated with MC3482 compared to vehicle. Data are presented as mean ± SEM from independent biological replicates, with each dot representing an individual donor. Statistical significance was determined using two-way ANOVA followed by Šídák’s multiple comparisons test. \**p* < 0.05, \*\**p* < 0.01, \*\*\**p* < 0.001.

### 3.4. SIRT5-dependent metabolic remodeling supports arginine biosynthesis in CD4^+^ T cells

To further evaluate the role of SIRT5 in NR-mediated arginine metabolism, targeted metabolomic profiling was performed in control and SIRT5-depleted CD4^+^ T cells. Interestingly, the most statistically robust metabolite affected by SIRT5 depletion was a distinct induction of the amino acid taurine, which is known to directly inhibit ASL function (Fig. 5A).[22] This finding further supported the concept that the arginine biosynthesis pathway was modified by SIRT5 levels. To examine this further control and SIRT5 knockdown (KD) CD4^+^ T cells were treated with NR or arginine and exposed to targeted metabolomic analysis. Interestingly, in control cells NR increased the steady-state levels of citrulline, arginine and ornithine, an amino acid intermediate downstream of arginine in the urea cycle (Fig. 5B and Fig. S3A). In contrast the levels of these urea cycle intermediates were not significantly changed in the absence of SIRT5 (Fig. 5B and Fig. S3B), supporting a requirement for SIRT5 in NR-mediated remodeling of the arginine biosynthetic pathway. The only other metabolite in this targeted analysis that was significantly induced by NR in the presence of SIRT5 was hypoxanthine, a purine derived metabolic intermediate (Fig. 5B). Quantitation of the normalized count values illustrates the effect of NR on the increase in arginine and citrulline abundance in control CD4⁺ T cells, whereas effect was abolished following SIRT5 depletion (Fig. 5C). Similarly, NR-induced increases in ornithine were attenuated in SIRT5-deficient cells (Fig. S3C), further supporting disruption of arginine biosynthetic flux downstream of ASL. To further evaluate the ASL node, arginosuccinate levels were quantified in control and SIRT5 KD cells following NR or arginine supplementation. Arginosuccinate accumulated in response to arginine supplementation in both control and SIRT5 KD cells, whereas the response to NR was comparatively attenuated in SIRT5-deficient cells (Fig. S3D). These findings suggest that SIRT5 promotes ASL-dependent flux from arginosuccinate toward arginine and fumarate production, thereby supporting arginine biosynthesis in human CD4⁺ T cells.

**Figure 5.**
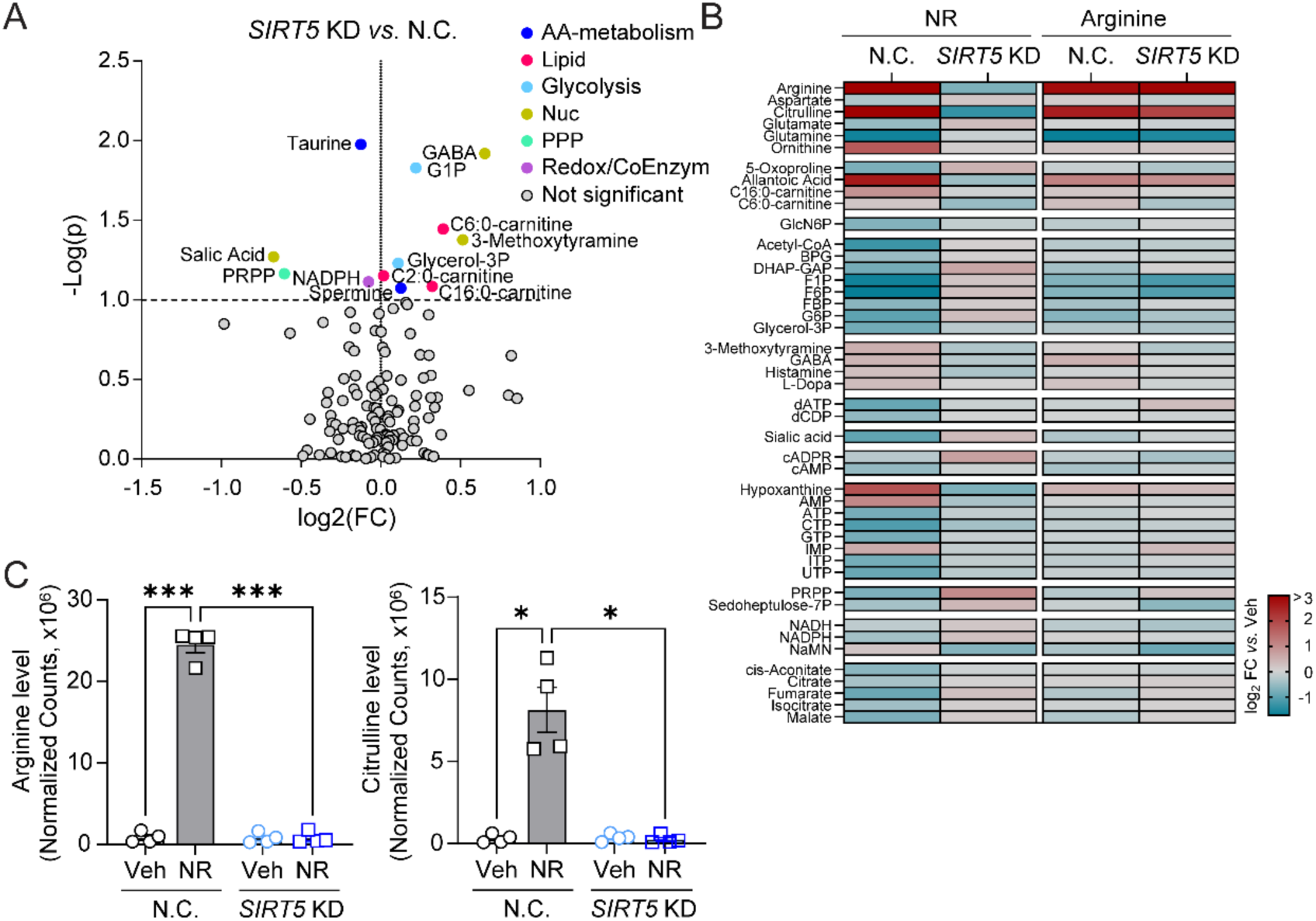
SIRT5-dependent metabolic remodeling supports arginine biosynthesis in CD4^+^ T cells. (**A**) Volcano plot of the pairwise comparison of SIRT5 knockdown (KD) *versus* non-targeting control (N.C.). Metabolites are plotted according to log₂ fold change and −log(*p*-value) and are color-coded by metabolic pathway. (**B**) Heatmap of metabolite abundance in N.C. and SIRT5 KD CD4+ T cells treated with NR or arginine. Data are shown as log2 fold change relative to vehicle-treated controls. Metabolomics data are provided in Supplementary Dataset 2. **(C-D)** Intracellular arginine (**C**) and citrulline (**D**) levels in N.C. and SIRT5 KD CD4⁺ T cells treated with vehicle or NR. Data are presented as normalized metabolite counts. Data are shown as mean ± SEM from independent biological replicates, with each symbol representing an individual donor. Statistical significance was determined using paired two-tailed Student’s *t*-test or two-way ANOVA with Šídák’s multiple comparisons test, as appropriate. \**p* < 0.05, \*\**p* < 0.001.

### 3.5. SIRT5 regulates ASL-dependent arginine metabolism and ASL acylation

To validate that SIRT5 contributed to arginine biosynthesis, we employed gain-of-function approaches in primary human CD4^+^ T cells. Overexpression of ASL significantly increased intracellular arginine levels, and this response was further augmented by NR supplementation (Fig. 6A). Similarly, SIRT5 overexpression increased arginine abundance under basal condition and following NR treatment (Fig. 6B). Co-expression of ASL and SIRT5 resulted in a greater increase in arginine levels compared to ASL expression alone, supporting a cooperative role for these proteins in the regulation of arginine biosynthesis (Fig. 6C). Consistent with this, NR supplementation further augmented arginine abundance in cells co-expressing ASL and SIRT5 (SFig. S4A). To determine whether SIRT5 directly interacts with ASL, HA-tagged SIRT5 was co-expressed with ASL-Flag in HEK293 cells. Expression of both proteins was confirmed by immunoblotting (Figures S4B-C). Reciprocal immunoprecipitation using anti-HA and anti-Flag antibodies demonstrated a robust interaction between SIRT5 and ASL, confirming their physical association (Fig. 6D). As SIRT5 functions primarily as a lysine deacylase, we next explored whether ASL displayed SIRT5-dependent changes in acylation. Flag immunoprecipitation followed by immunoblotting demonstrated SIRT5-dependent reductions in ASL glutarylation, supporting a role for SIRT5 in regulating ASL acylation (Fig. 6E). Mutation of candidate lysine residues to arginine altered ASL acylation patterns, with K69R, K288R, and double mutant (DM) constructs exhibiting distinct glutarylation profiles compared with wild-type ASL (Fig. 6E and Fig. S4E). Quantification of glutaryl-lysine signal confirmed significant differences among the mutant constructs, supporting an important role for K69 to SIRT5-dependent ASL deglutarylation (Fig. 6F). To further evaluate whether additional lysine acylation species contribute to ASL regulation, ASL acetylation was assessed following HA immunoprecipitation. In contrast to glutarylation, malonyl- and acetyl-lysine modification showed relatively modest changes across the ASL mutants (Fig. S4E), suggesting that SIRT5-dependent regulation of ASL is preferentially associated with glutaryl-lysine modifications. Collectively, these findings identify ASL as a SIRT5-interacting protein and support a model in which SIRT5 regulates arginine biosynthesis through modulation of ASL glutarylation. Of note, in CD4^+^ T cells, antibodies could not appreciate succinyl-modifications but cannot ascertain if this is due to their low abundance or due to poor antibody affinity (data not shown).

**Figure 6.**
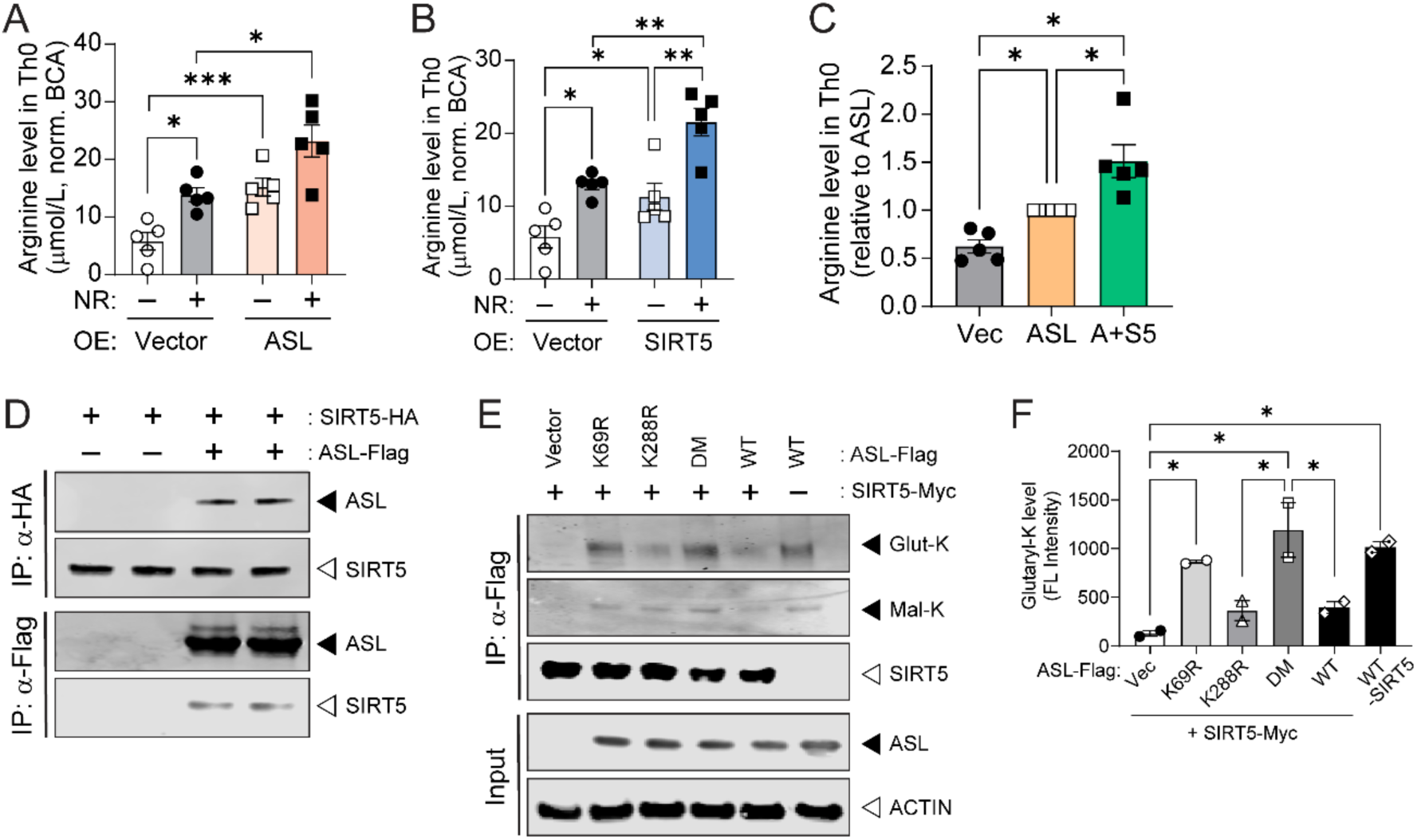
SIRT5 regulates ASL-dependent arginine metabolism and ASL acylation. (**A-B**) Intracellular arginine levels in CD4⁺ T cells overexpressing ASL (A) or SIRT5 (B) in the presence or absence of NR. (**C**) Intracellular arginine levels in Th0 cells overexpressing ASL alone or in combination with SIRT5. (**D**) Reciprocal co-immunoprecipitation analysis demonstrating interaction between SIRT5 and ASL. HA-IP and Flag-IP confirm the association between SIRT5-HA and ASL-Flag. (**E**) ASL glutarylation assessed by Flag-IP followed by immunoblotting for glutaryl-lysine (Glut-K) in wild-type (WT),K69R, K288R, and double mutant (DM) ASL constructs in the presence of SIRT5. (**F**) Quantification of ASL glutarylation shown in (E). Data are presented as mean ± SEM from independent biological replicates, with each dot representing an individual donor. Statistical significance was determined using one-way or two-way ANOVA with Šídák’s multiple comparisons test, as appropriate. \**p* < 0.05, \*\**p* < 0.01, \*\*\**p* < 0.001.

## 4. Discussion

In this study we identify the sirtuin SIRT5, is an additional regulatory node contributing to NAD^+^-mediated dampening of Th1 and Th17 immune responsiveness. Here, the NAD^+^-dependent SIRT5, appears via the posttranslational deacylation of arginosuccinate lyase, to enhance arginine biosynthesis. This places the role of NR-activated SIRT5 upstream of the redox signaling and transcriptional control in attenuating CD4^+^ immune responsiveness.[9]

SIRT5 is a distinct sirtuin in that is highly NAD^+^-dependent,[23] possesses weak deacetylase activity, and preferentially enables deacylase activity to catalyze the removal of succinyl, malonyl and glutaryl intermediates from protein lysine residues. Although these distinct effects have not been comprehensively defined, initial studies implicate that these distinct deacylase activities may be enzyme/metabolic pathway distinct. SIRT5 mediated desuccinylation appears to function in anaplerotic and mitochondrial oxidative metabolism.[24, 25] The role of SIRT5 in demalonylation appears to preferentially target glycolysis and the gluconeogenesis pathways,[25] and its role in deglutarylation in amino acid metabolism.[26] SIRT5 has also been shown to modulate the urea cycle via the modification and activation of carbamoyl phosphate synthetase 1.[18] This study further supports a role for SIRT5 in the urea cycle, and more specifically in modification of the functional intersection enzyme, arginosuccinate lyase (ASL), which is shared between the urea cycle and arginine biosynthesis. Targeted metabolomic profiling further supported a role for SIRT5 in regulating the ASL node within the arginine biosynthetic pathway. SIRT5 depletion attenuated NR-induced increases in citrulline, arginine, and ornithine, while promoting accumulation of arginosuccinate. Together, these findings are consistent with impaired ASL-dependent flux toward arginine and fumarate production in the absence of SIRT5. In parallel, biochemical studies identified ASL as a SIRT5-interacting protein and demonstrated that ASL glutarylation was the predominant acyl modification regulated by SIRT5. Although we could not comprehensively assess ASL succinylation, these findings support a role for SIRT5-mediated deglutarylation in the regulation of ASL-dependent arginine metabolism.

In the pantheon of immunometabolism, arginine is well established in myeloid cell biology with pro- and anti-inflammatory effects as a substrate for nitric oxide and in polyamine biology respectively.[27] Arginine and its byproduct fumarate from arginosuccinate catabolism have been shown to regulate CD4^+^ T cell biology via activation of the NRF2 transcription factor.[9] This study expands on that work by demonstrating that ASL-dependent arginine metabolism is regulated by NAD^+^-dependent SIRT5 signaling and further advances our understanding of how nutrient-sensing integrates NAD^+^ metabolism with the regulation of Th1 and Th17 immune responsiveness.

In myeloid cells, SIRT5 has been directly shown to rescue the inflammatory response in endotoxin-tolerant macrophages[28] and to orchestrate proinflammatory polarization of resident macrophages,[29] both via deacylation effects. Conversely, SIRT 5 blunts macrophage IL-1β in a murine colitis model,[30] and SIRT5 levels are inversely correlated with human rheumatoid arthritis, and exogenous SIRT5 attenuated LPS-induced inflammation in primary human monocytes.[31] Together, these studies illustrate the pro-and anti-inflammatory effects of SIRT5. However, the role of arginine biochemistry in myeloid cell immunoregulation has not explored been explored. Whether context specific effects of SIRT5 on arginine metabolism contribute to these divergent myeloid cell inflammatory programs remains an important question for further investigation.

The putative overlap between myeloid and lymphoid biology in response to NR administration is further postulated by the identification of hypoxanthine as a NR-regulated metabolite in CD4^+^ T cells. Previously NR has been found to increase purine metabolite levels in primary human monocytes, and these metabolites, including hypoxanthine and inosine mimic the effect of NR in blunting monocytic type I interferon production.[10] Interestingly, in that study, the partial knockdown of SIRT5, did not ameliorate the anti-inflammatory effect of NR.[10] Here too, the role of purine metabolism, specifically hypoxanthine which was marked increased by NR, on CD4^+^ T cell biology and its interaction with SIRT5 will require further study.

In conclusion, this study expands our understanding of the CD4^+^ T cell immunoregulatory effects of NR supplementation to include activation of the NAD^+^-dependent sirtuin SIRT5. Furthermore, these findings place the activation of SIRT5 upstream of arginine biosynthesis via its effect on arginosuccinate lyase, probably via the deglutarylation of this enzyme, and broaden the role of sirtuin biology in adaptive immune cell regulation.

## Acknowledgments

We thank Dr. Shi-Min Zhao (Fudan University, Shanghai, China) for generously providing the human ASL expression plasmid used in this study. We also acknowledge the support of institutional core facilities for flow cytometry and metabolic analyses. This work was supported by the NIH Intramural Research Program of the National Heart, Lung, and Blood Institute (NHLBI) (to M.N.S.).

## Author contributions

Conceptualization, K.H. and M.N.S.; Methodology, K.H., R.J.K., K.W., A.C.R., A.M.M., T.C.R., R.S., and A.K.G.; Investigation, K.H., R.J.K., K.W., A.C.R., A.M.M., T.C.R., R.S., and A.K.G.; Formal Analysis, K.H., R.J.K., K.W., A.C.R., A.M.M., T.C.R., R.S., A.K.G., N.T.B., and B.S.; Resources, N.T.B., B.S., and M.N.S.; Writing – Original Draft, K.H.; Writing – Review & Editing, K.H., R.J.K., K.W., A.C.R., A.M.M., T.C.R., R.S., A.K.G., N.T.B., B.S., and M.N.S.; Supervision, M.N.S.; Funding Acquisition, M.N.S.

## INCLUSION AND DIVERSITY

We support inclusive, diverse, and equitable conduct of research.

## Data availability

The datasets supporting the conclusions of this article are available from the corresponding author upon reasonable request.

