## Supplementary material for "SIRT5-dependent regulation of ASL controls arginine metabolism and T cell function": Suppl Fig + table

### SUPPLEMENTAL MATERIALS

- **Supplemental Figures and Legends (4)**
- **Supplemental Table (1)**
- **Supplemental Datasets (2) - Uploaded separately as Excel files**

**Supplemental Dataset 1:** Integrated multi-omics dataset (lipidomics, metabolomics, and RNA-seq), related to Figure 1

**Supplemental Dataset 2:** Metabolomics dataset of SIRT5-dependent metabolic remodeling, related to Figure 5

### Supplemental Figures and Legends

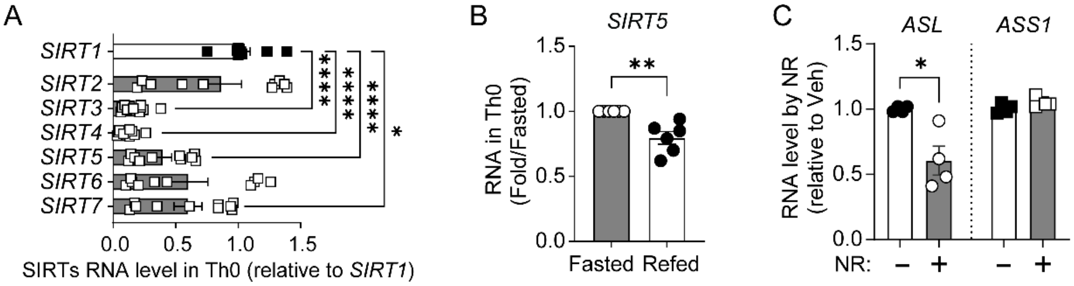


**Supplementary Figure 1. Expression of sirtuins and arginine pathway genes in CD4$^+$ T cells.** (**A**) Relative RNA expression of sirtuin family members (*SIRT1-7*) in CD4$^+$ T cells under Th0 conditions, normalized to *SIRT1*. (**B**) Relative expression of *SIRT5* in CD4$^+$ T cells under fasting and refeeding conditions. (**C**) Relative RNA expression of *ASL* and *ASS1* in CD4$^+$ T cells treated with NR compared to vehicle. Data are presented as mean ± SEM from independent biological replicates, with each dot representing an individual donor. Statistical significance was determined using one-way ANOVA with Šídák's multiple comparisons test (A) or paired two-tailed Student's *t*-test (B-C). *$p < 0.05$, **$p < 0.01$, ***$p < 0.001$.

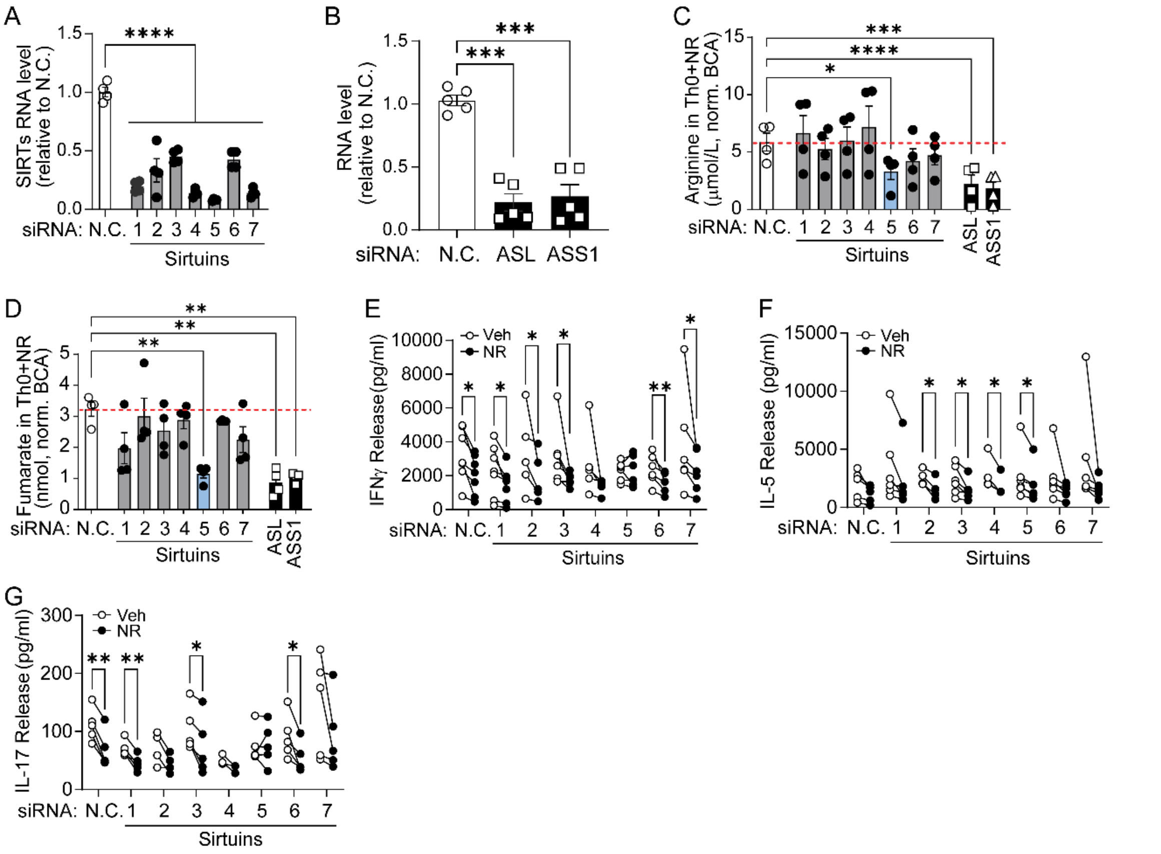


**Supplementary Figure 2. Sirtuin screening identifies SIRT5 as a regulator of arginine metabolism in CD4⁺ T cells.** (**A**) Relative RNA expression of sirtuins following siRNA-mediated knockdown, normalized to non-targeting control (N.C.). (**B**) Relative RNA expression of *ASL* and *ASS1* following siRNA-mediated knockdown. (**C-D**) Intracellular arginine and fumarate levels in $CD4^+$ T cells transfected with siRNAs targeting individual sirtuins (SIRT1-7), ASL, or ASS1 in the presence of NR. (**E-G**) Cytokine production (IFNγ, IL-5, and IL-17) in $CD4^+$ T cells transfected with siRNAs targeting individual sirtuins and treated with vehicle or NR. Data are presented as mean ± SEM from independent biological replicates, with each dot representing an individual donor. Statistical significance was determined using one-way ANOVA with Šídák's multiple comparisons test (A-D) or two-way ANOVA with Šídák's multiple comparisons test (E-G). $*p < 0.05$, $**p < 0.01$, $***p < 0.001$.

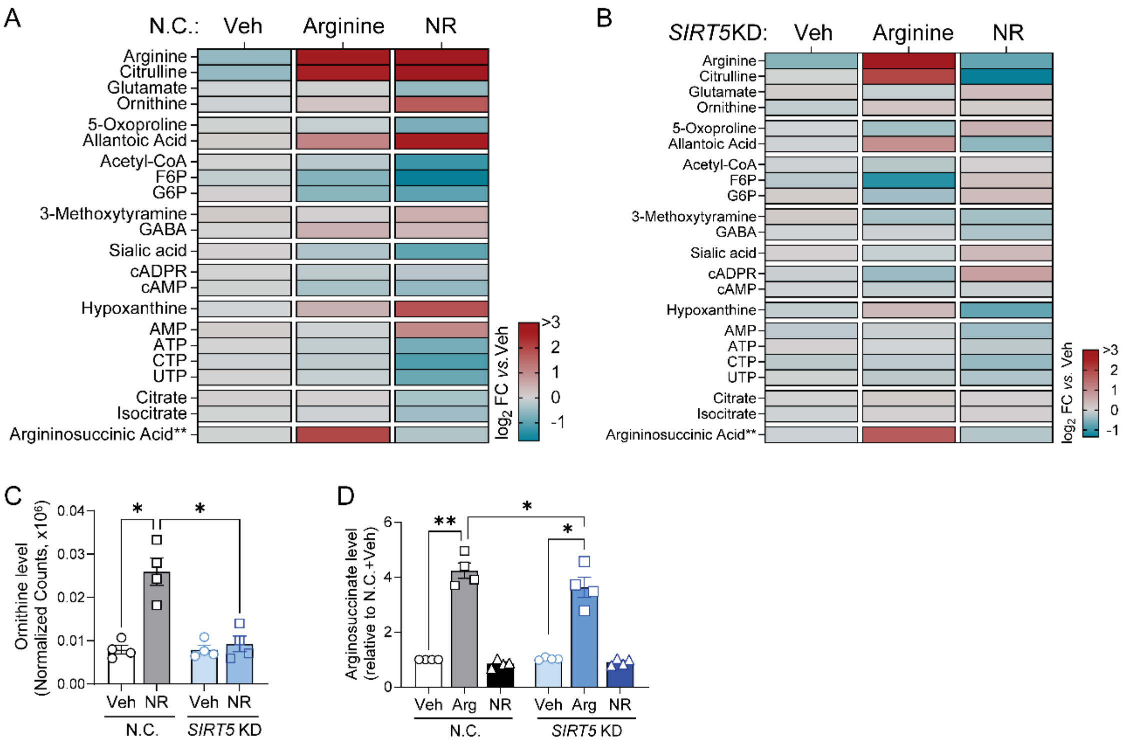


**Supplementary Figure 3. Metabolomic responses to NR and arginine in control and SIRT5-dependent CD4$^+$ T cells. (A-B)** Heatmaps showing metabolite abundance in non-targeting control (N.C., **A**) and SIRT5 knockdown (KD, **B**) CD4$^+$ T cells treated with vehicle, arginine, or NR. Data are presented as $\log_2$ fold change relative to vehicle-treated controls. Data are presented as $\log_2$ fold change relative to vehicle-treated controls. **(C)** Intracellular ornithine levels in N.C. and SIRT5 KD CD4$^+$ T cells treated with vehicle or NR. **(D)** Relative arginosuccinate levels in N.C. and SIRT5 KD CD4$^+$ T cells treated with vehicle, arginine, or NR. Arginosuccinate abundance was normalized to the N.C. vehicle condition. Data are presented as mean ± SEM from independent biological replicates, with each symbol representing an individual donor. Statistical significance was determined using paired two-tailed Student's *t*-test or two-way ANOVA with Šídák's multiple comparisons test, as appropriate. $*p < 0.05$, $**p < 0.01$.

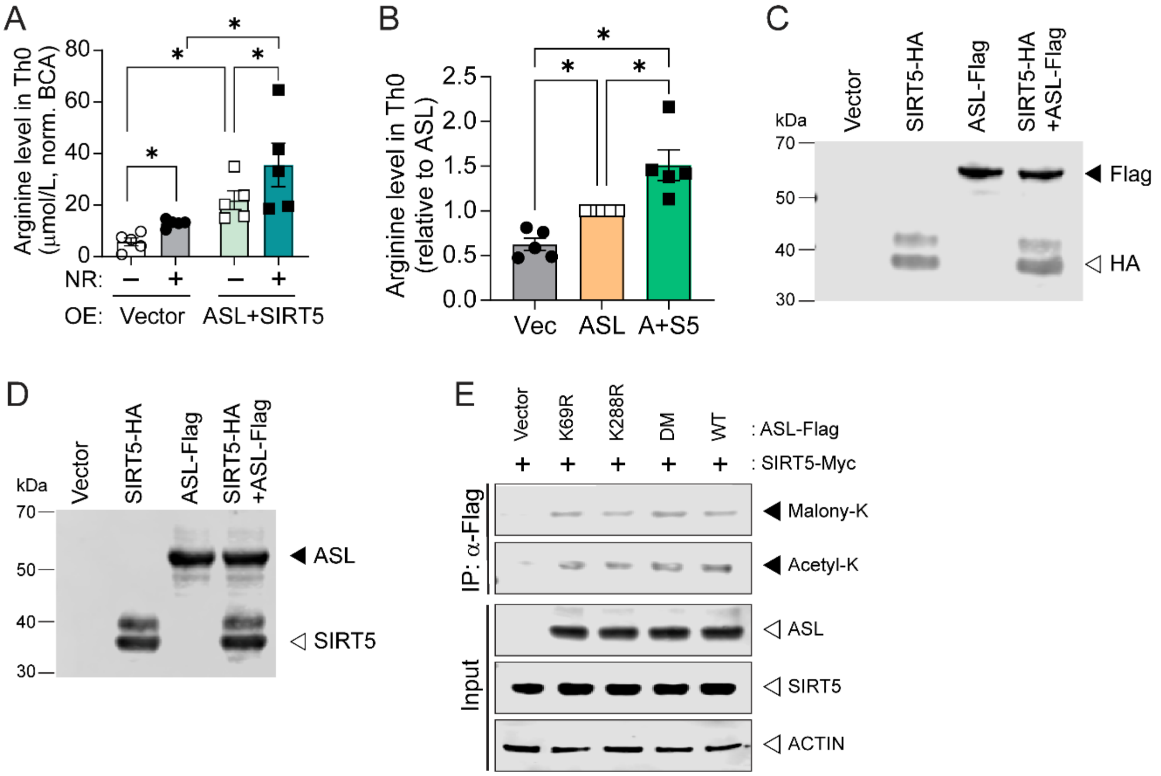


**Supplementary Figure 4. Overexpression of SIRT5 and ASL regulates arginine metabolism and ASL acylation.** (**A**) Intracellular arginine levels in $CD4^+$ T cells overexpressing ASL and SIRT5 in the presence or absence of NR. (**B-C**) Immunoblot analysis confirming expression of SIRT5-HA and ASL-Flag constructs used for co-immunoprecipitation studies. (**D**) ASL malonylation (Malonyl-K) and acetylation (Acetyl-K) assessed by Flag-IP and followed by immunoblotting in wild-type (WT), K69R, K288R, and double mutant (DM) in the presence of SIRT5. Data in (A-B) are presented as mean ± SEM from independent biological replicates. Statistical significance was determined using two-way ANOVA followed by Šídák's multiple comparisons test. *$p < 0.05$.

**Supplemental Table 1.** Pre-made primer sets used for quantitative real-time RT-PCR.

| Oligonucleotides | SOURCE | IDENTIFIER |
|---|---|---|
| *SIRT1* | Qiagen | QT00051261 |
| *SIRT2* | Qiagen | QT00069531 |
| *SIRT3* | Qiagen | QT00091490 |
| *SIRT4* | Qiagen | QT00202503 |
| *SIRT5* | Qiagen | QT00047537 |
| *SIRT6* | Qiagen | QT00056812 |
| *SIRT7* | Qiagen | QT01020663 |
| *ASL* | Qiagen | QT00054579 |
| *ASS1* | Qiagen | QT00036456 |
| *RRN18S* | Qiagen | QT00199367 |
| *b-ACTIN* | Qiagen | QT00095431 |
